# The effects of mechanical strain on mouse eye lens capsule and cellular microstructure

**DOI:** 10.1101/247502

**Authors:** Justin Parreno, Catherine Cheng, Roberta B. Nowak, Velia M. Fowler

## Abstract

The understanding of multiscale load transfer within complex soft tissues is incomplete. The eye lens is ideal for multiscale tissue mechanics studies as its principal function is to fine focus light at different distances onto the retina via mechanical shape changes. The biomechanical function, resiliency, and intricate microstructure of the lens make it an excellent non-connective soft tissue model. We hypothesized that compressive strain applied onto whole lens tissue leads to deformation of specific microstructures and that this deformation is reversible following removal of load. For this examination, mouse lenses were compressed by sequential application of increasing load. Using confocal microscopy and quantitative image analysis, we determined that axial strain ≥10% reduces capsule thickness, expands epithelial cell area, and separates fiber cell tips at the anterior region of the lenses. At the equatorial region, strain ≥6% increases fiber cell widths. The effects of strain on lens epithelial cell area, capsule thickness, and equatorial fiber cell widths are reversible following the release of lenses from strain. However, although fiber cell tip separation following the removal of low loads is reversible, the separation becomes irreversible with application of higher loads. This irreversible separation between fiber cell tips leads to incomplete bulk lens resiliency. The lens is an accessible biomechanical model system that provides new insights on multiscale transfer of loads in soft tissues.

## Introduction

An understanding of the multiscale relationship between macro- and micro- level mechanics is essential for determining how tissue mechanical properties emerge from specific tissue microstructures (Dumont and Prakash, 2014). The multiscale transfer of load in soft tissues has been previously characterized in studies that use isolated portions of connective tissues (such as tendon, meniscus, and annulus fibrosis) (Bruehlmann *et al*., 2004; Upton *et al*., 2008; Desrochers and Duncan, 2010; Han *et al*., 2013). In these connective tissues, the proportion of strain applied at the tissue level is largely-to-completely attenuated by the extracellular matrix (ECM), prior to reaching the small number of cells embedded within a large amount of ECM (Bruehlmann *et al*., 2004; Han *et al*., 2013). For a more complete understanding of microscale load transfer in tissues, studies that examine how loads are distributed within more complex tissues that have a vastly different microstructural cellular composition and organization than connective tissues are needed.

The eye lens serves as an ideal model to develop new insights into the multiscale transfer of mechanical loads in tissues for several reasons. The lens is a soft, transparent tissue in the anterior chamber of the eye that changes shape to fine focus light from different distances onto the retina to generate a clear image. Thus, the optical function of the lens is closely tied to its mechanical function. The biomechanical stiffness of soft-tissues (Candiello *et al*., 2010; Tsujii *et al*., 2017; Lampi and Reinhart-King, 2018), including the lens (Baradia *et al*., 2010; Hozic *et al*., 2012; Cheng *et al*., 2016a) increase with age. Increased biomechanical tissue stiffness in the aging lens diminishes its ability to change shape, resulting in presbyopia and the need for reading glasses (Heys *et al*., 2004). Therefore, determining the role of lens microstructure during bulk lens shape change will provide a foundation for understanding lens pathology as well as the general relationship between microstructure and age-related stiffness of soft-tissues. Another property that could provide additional insights into multiscale load transfer is lens resiliency. Release from tensional strain, *in vivo*, leads to lens rounding. While the effect of mechanical unloading has recently been shown in cells (Bonakdar *et al*., 2016), the effect of unloading on microstructures within a tissue system has not been determined. Thus, the lens also provides an opportunity to investigate microstructure during release from various magnitudes of mechanical loads.

The lens serves as an important model of multiscale load transfer as it is a predominantly cellular tissue, with a microstructural composition and organization dramatically different from connective soft-tissues (Upton *et al*., 2008; Desrochers and Duncan, 2010; Han *et al*., 2013; Szczesny and Elliott, 2014). The only extracellular matrix present in the lens is a thin basement membrane, known as the lens capsule. The capsule, which is rich in type IV collagen, surrounds the entire organ (Lovicu and Robinson, 2004). Inward from the capsule are the lens cells, which have intricate shapes and are highly organized (Wanko and Gavin, 1959; Cheng *et al*., 2017b). A monolayer of epithelial cells covers the anterior hemisphere of the lens and is attached to the capsule. Lifelong lens growth is facilitated by continuous proliferation and differentiation of equatorial epithelial cells into newly formed fiber cells. Fiber cells migrate and elongate into thin, ribbon-like cells extending from the anterior to posterior pole, and new layers of fiber cells are added onto older generations of fiber cells in concentric shells. At the poles, fiber cell tips from the opposing sides of the lens meet to form the anterior and posterior suture cell-cell junctions (Kuszak *et al*., 1984; Kuszak *et al*., 2006; Kuszak and Zoltoski, 2006). Thus, determination of multiscale load transfer in the lens will aid in elucidating tissue responses to load for complex and predominantly cellular tissues.

In this study, we examined the hypothesis that compressive strain leads to reversible deformation of lens microstructural dimensions. We developed new quantitative methodology to examine lens capsule thickness, epithelial cell area, fiber cell widths, and fiber tip organization during, as well as recovery from, compression in live mouse lenses. This analysis has allowed us to characterize multiscale load transfer in the lens. We show that, in contrast to connective tissues (Bruehlmann *et al*., 2004; Upton *et al*., 2008; Desrochers and Duncan, 2010; Han *et al*., 2013), load not only transferred to the lens matrix (capsule), but also throughout the lens epithelial and fiber cells. We also determined that whole lens resiliency was incomplete following the release from high loads and that this corresponded to incomplete recovery of fiber cell tip organization. Other microstructural features, however, such as capsule thickness and equatorial fiber widths, were completely recovered following release from high loads, suggesting that the inability of tissues to recover from load could be due to failure of specific microstructures. The lens model provides a robust system to develop insights into multiscale load transfer onto cells through a thin matrix that will result in a better understanding of how biomechanical tissue properties emerge from specific tissue microstructures.

## Results

### tdTomato labeling of mouse lenses allows live visualization of the lens anterior

We utilized mice expressing tandem dimer-Tomato (B6.129(Cg)-Gt(ROSA) (hereafter referred to as tdTomato) to visualize plasma membranes of the cells in live lenses. These mice have been genetically engineered to express tdTomato protein fused to connexin 43 and driven by the chicken β-actin core promoter with a CMV enhancer (the tdTomato-connexin 43 communication is disabled) (Campbell *et al*., 2002; Muzumdar *et al*., 2007). This results in membrane targeting of tdTomato and allows for visualization of cell morphology by confocal fluorescence microscopy.

Although expression of tdTomato in these mice is ubiquitous, the expression can be variable depending on tissue type (Muzumdar *et al*., 2007). Therefore, we examined whether the tdTomato expression was present at epithelial and fiber cell plasma membranes in the anterior region of live mouse lenses (as depicted in Figure 1A). We observed that tdTomato labeling is bright and stable and can be easily observed in lens cells as deep as 150μm from the capsule surface (Figure 1B). Additionally, the characteristic cobble stone morphology of the anterior epithelial cell monolayer and the fiber cell tips at the Y-shaped sutures of mouse lenses (Kuszak *et al*., 1984; Cheng *et al*., 2017b) are apparent with endogenous tdTomato labeling (Figure 1C).

**Figure 1.**
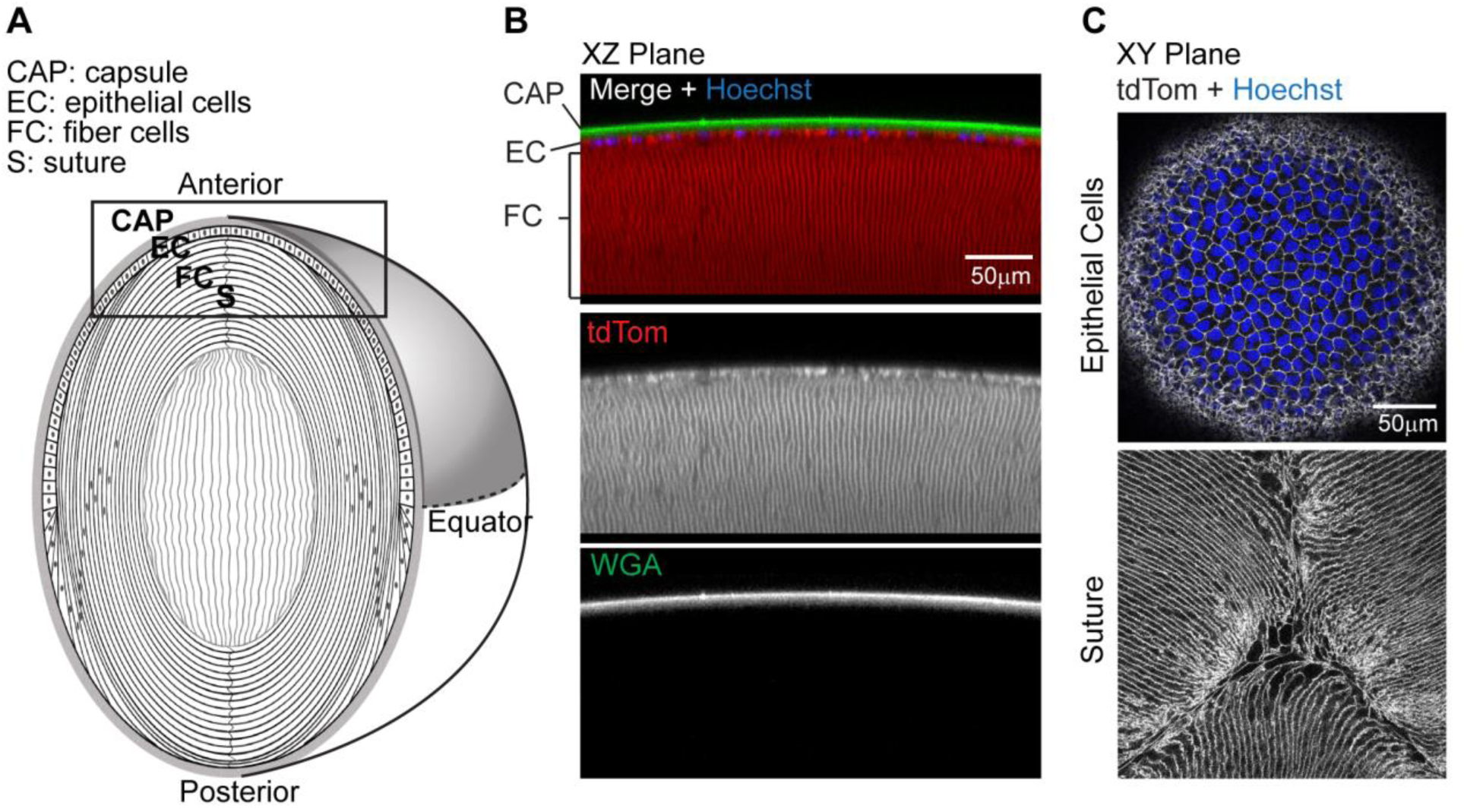
tdTomato-labeled membranes in mouse lenses allow live cell confocal imaging. (A) Diagram of lens anatomy showing lens capsule (CAP), epithelial cells (EC), fiber cells (FC) and suture (S). The lens anterior region is boxed and the lens equator is indicated by the dashed horizontal line. (B) Sagittal optical section (XZ plane view) of reconstructed confocal Z-stacks of the lens anterior showing epithelial cells and fiber cell membranes labeled with tdTomato (red), lens capsule stained with WGA (green), and epithelial cell nuclei stained with Hoechst (blue). (C) Single optical sections (XY plane view) of tdTomato-labeled membranes of lens epithelial and fiber cells (gray scale), showing Hoechst-stained nuclei (blue) in the epithelium (top panel), and the anterior fiber cells meeting at the suture (bottom panel). Scale bars, 50μm.

### Biomechanical properties of wild-type and tdTomato lenses

TdTomato is targeted to the plasma membrane and since cell membrane properties contribute to cell mechanical properties (Heidemann and Wirtz, 2004), we next sought to determine whether tdTomato membrane labeling alters the bulk biomechanical properties of the lens. To perform this analysis, we sequentially applied increasing axial load in the form of glass coverslips onto tdTomato (Figure 2A) or wild-type C57BL/6 mouse lenses and determined axial and equatorial strain from measurements of the lens axial and equatorial diameters, respectively. This method allows for precise and reproducible measurement of lens strain (Baradia *et al*., 2010; Gokhin *et al*., 2012; Cheng *et al*., 2016a). We determined that at any given load tested, there are no differences (p > 0.05) in axial strain between wild-type and tdTomato lenses (Figure 2B). For subsequent experiments, we utilize a range of strains by applying either 1, 2, 5, or 10 coverslips onto lenses. These loads result in axial strains of 10%, 14%, 23%, and 29% in mouse lenses, respectively.

**Figure 2.**
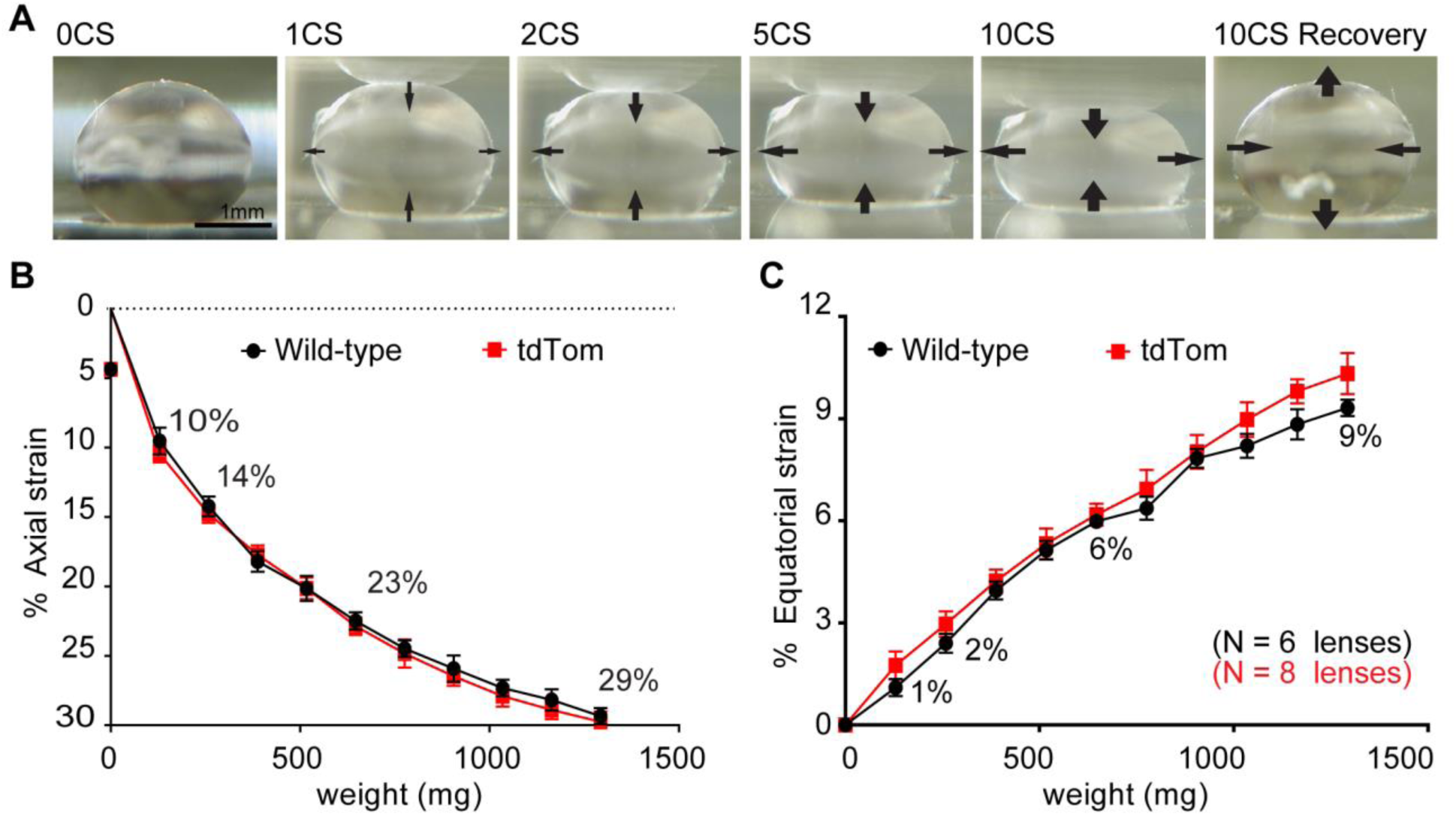
The effect of coverslip compression on mouse lens axial and equatorial strain. Sagittal view images of tdTomato mouse lens compressed by the indicated number of coverslips (CS) and following the removal of 10 coverslips (10CS recovery). Arrows on images indicate direction of lens shape changes at the anterior and posterior, and at the equator. Plots of (B) axial and (C) equatorial strains as a function of coverslip weight for wild-type and tdTomato mouse lenses. The application of 1, 2, 5, and 10 coverslips (129.3 mg per coverslip) provided a range of strains that were used for the remainder of this study. Numbers are % strain calculated for the indicated numbers of coverslips.

Lenses behave according to the Poisson effect such that axial compression also results in equatorial expansion (Bailey *et al*., 2010; Gokhin *et al*., 2012; Cheng *et al*., 2016a). Measurements of equatorial strain of the lens during compression reveal that, at any given load, the magnitude of axial strain (Figure 2B) is greater than equatorial strain (Figure 2C). Similar to axial strain, at any given load tested, there are no differences (p > 0.05) in equatorial strain between wild-type and tdTomato lenses (Figure 2C). The application of 1, 2, 5, and 10 coverslip loads results in equatorial strains of 1, 2, 6, and 9% in mouse lenses, respectively.

### Bulk lens dimensions recover only partially at higher strains

To examine if the amount of strain affected recovery to the original lens shape after release from compression, we determined if there were differences between pre- and post-axial and equatorial lens diameters and calculated the percent recovery. Initial studies indicate no differences in recovery of axial diameter between tdTomato and wild-type mouse lenses (Figure S1), therefore we pooled data from tdTomato and wild-type lenses. Release from 10% axial strain results in complete axial diameter recovery as there is no significant difference (p > 0.05) between pre- and post-strain axial lens diameters (Figure 3A). Release from axial strains ≥14%, however, results in only partial recovery of axial diameter as the post-strain axial diameter is significantly less compared to the pre-strain axial diameter. Axial strains of 14%, 23%, and 29% led to 98.0±1.0% (p = 0.01), 96.5±2.8% (p = 0.03), and 95.9±1.1% (p < 0.001) recovery of axial diameter, respectively (Figure 3B).

**Figure 3.**
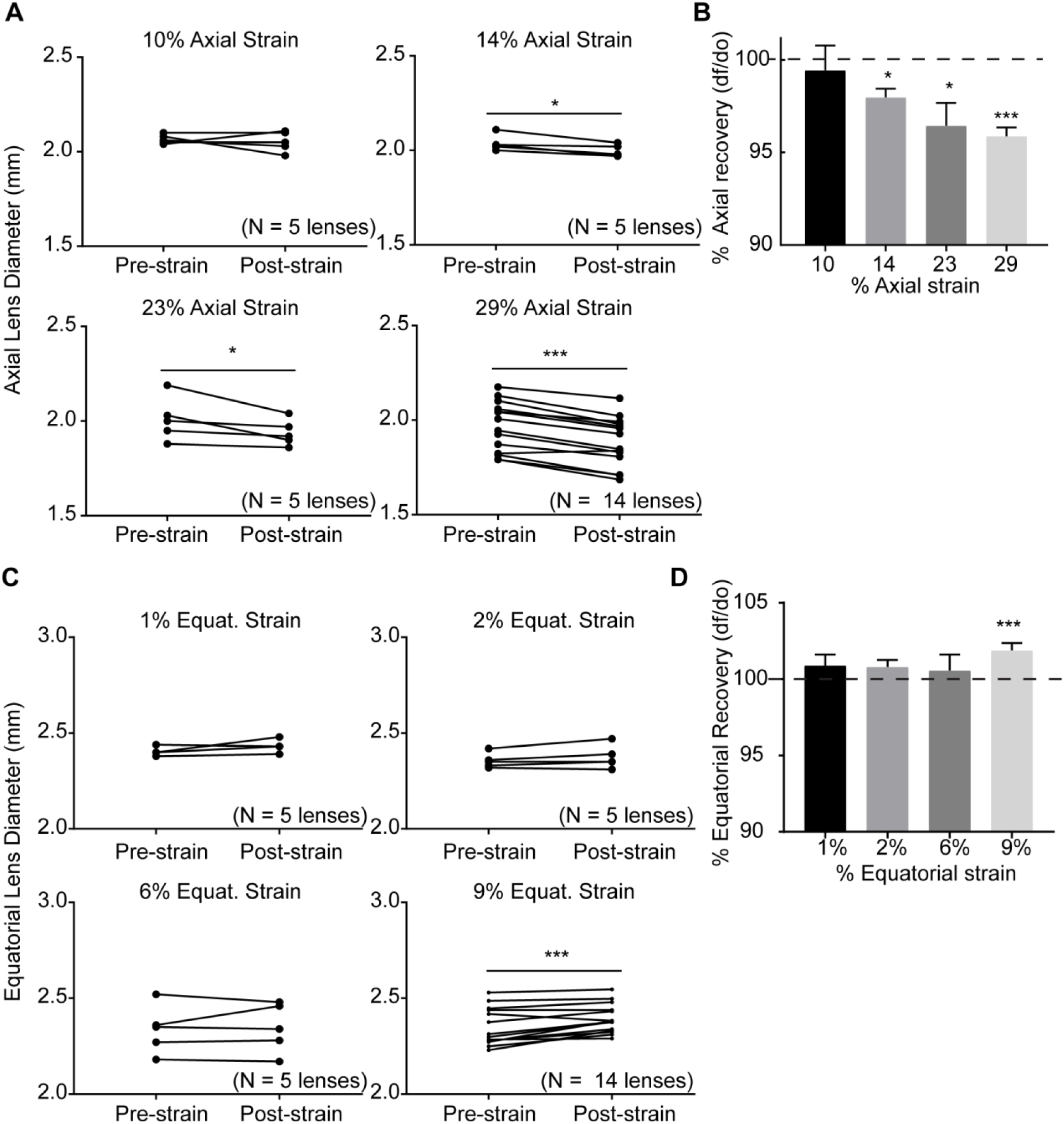
Recovery of bulk lens axial and equatorial dimensions following release from strain. (A) Plots of axial diameters of lenses pre- and post-strain. (B) Bar graphs showing that axial diameter recovery (percentage of post- to pre-axial diameter) progressively decreased with increasing strain. (C) Plots of equatorial diameters of lenses pre- and post-strain. (D) Bar graphs show that equatorial diameter recovery (percentage of post- to pre-equatorial diameter) is incomplete at 9% equatorial strain. *, p < 0.05; **, p < 0.01; ***, p < 0.001.

Similar to recovery of lens axial diameter, recovery of lens equatorial diameter was complete at lower equatorial strains. Release from equatorial strains ≤6% results in complete recovery of equatorial diameter as there was no significant difference (p > 0.05) between pre- and post- equatorial diameters. Release from 9% equatorial strain, however, results in partial recovery of equatorial diameter, as post-strain diameter is significantly greater (p < 0.001) than pre-strain diameter (Figure 3C), by 1.9% (Figure 3D).

Since recovery from high strains was incomplete, this suggests that increasing strain may lead to progressive tissue damage. Our data shows that compression with 10% axial strain in mouse lenses does not cause irreversible damage to bulk lens shape. Ten percent axial strain is within the range of physiological primate lens shape changes during accommodation calculated from *in vivo* measurements (Jackson, 1907; Doyle *et al*., 2013; Marussich *et al*., 2015). Thus, for the rest of this study, we focused primarily on the microstructural changes that occur at 10% axial strain. To provide additional insights into the microstructural changes that occur at high loads, we also examined the effects of higher levels of strain.

### Lens flattening caused by strain results in a reversible decrease in capsule thickness

Isolated human lens capsules are highly elastic and can stretch up to ~108% (Fisher, 1969b; Krag *et al*., 1997), even more so than other types of collagenous connective tissue (Oxlund and Andreassen, 1980). This propensity to undergo strain has led to the speculation that capsules stretch during lens flattening (Fisher, 1969a). To examine the effect of axial strain on capsule deformation, we measured capsule thickness in the anterior region of lenses (as depicted in Figure 4A) prior to, during, and after the release from strain. For capsule visualization, live lens capsules were stained with a wheat germ agglutinin (WGA) fluorescent conjugate. In 3D reconstructed images, we determined that WGA staining is present throughout the lens capsule and was most intense at the top surface (Figure 4B). There is less staining for WGA in the underlying epithelial cells. In contrast to WGA staining, tdTomato labels the underlying epithelial cell membranes but not the capsule. These mutually exclusive labeling properties of WGA for the capsule and tdTomato for epithelial cells allow for precise measurement of capsule thickness by line scan analysis. Fluorescent line scan profiles reveal a WGA peak corresponding to the intense top surface staining of the capsule and a tdTomato peak corresponding to the basal surface of the epithelial membranes adjacent to the bottom surface of the capsule. The difference in distance between the two peaks is therefore capsule thickness.

**Figure 4.**
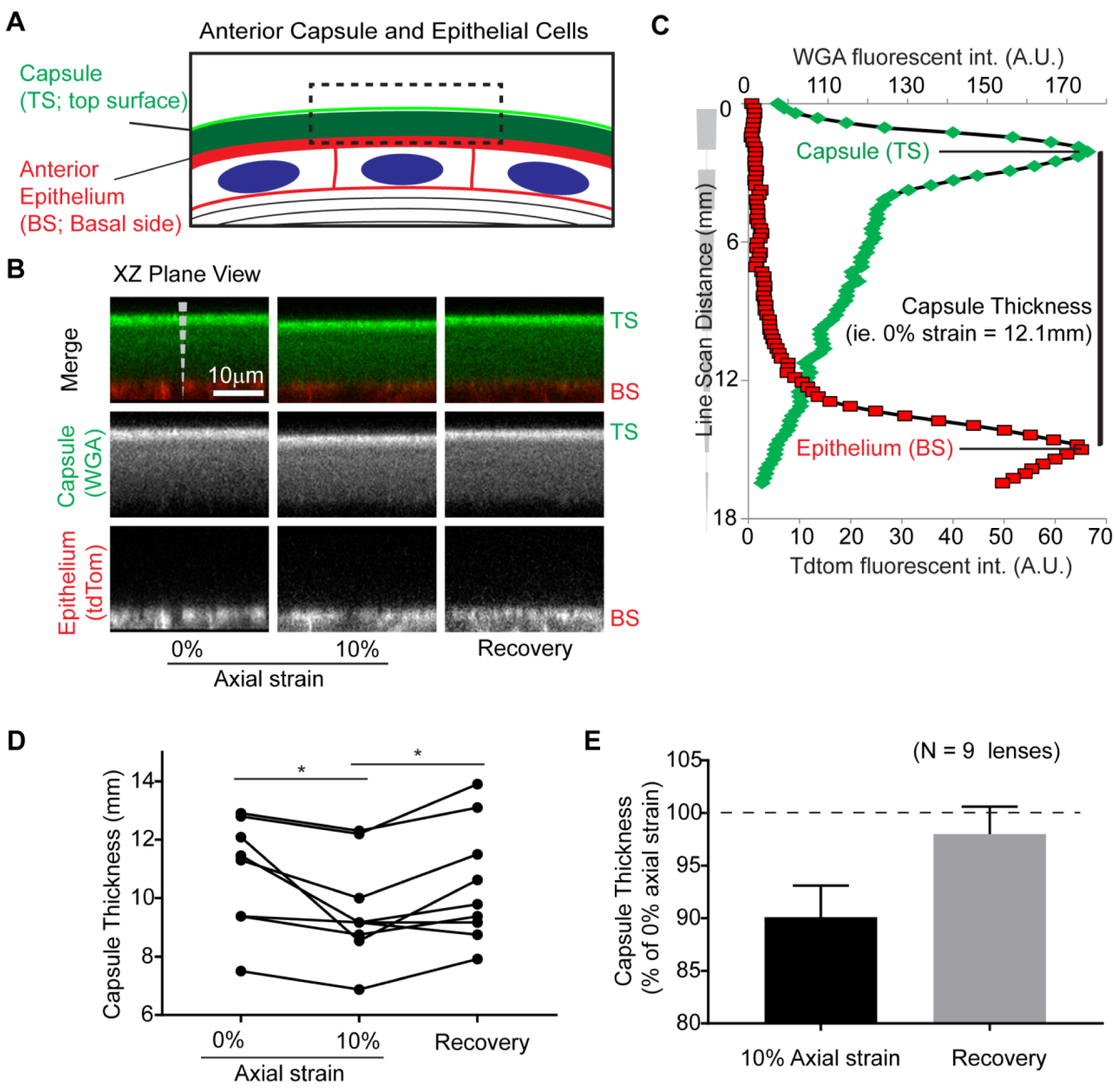
Lens axial strain leads to a reversible decrease in anterior capsule thickness. (A) Diagram depicting anterior capsule and epithelial cells. (B) Images of WGA-stained capsule (green) and tdTomato (tdTom)-labeled underlying epithelial cells (red) in a sagittal optical section (XZ plane view) from 3D reconstructions of confocal Z-stacks. Images from lenses at 0% and 10% strain, and following release from strain (Recovery). Line scans, indicated by dashed vertical white lines, with the taper of dashed line indicating the direction of line scan. Scale bar, 10μm. (C) Example of corresponding line scan data from images in B, demonstrating a single WGA (green) peak that corresponds to the top surface (TS) of capsule and a tdTomato (red) peak that corresponds to the basal side (BS) of epithelial cells adjacent to the capsule. Capsule thickness was determined by the distance between green and red peaks as indicated. (D) Repeated measures dot-plots of capsule thickness from unstrained (0%), 10% strained, and recovered lenses. (E) Average change in capsule thickness calculated from data in D, normalized to capsule thickness at 0% axial strain. *, p < 0.05; **, p < 0.01; ***, p < 0.001

We observed that anterior capsule thickness is variable between individual lenses (Figure 4D), in keeping with previous findings for C57BL/6 wild-type mice of similar ages (Danysh *et al*., 2008). Due to this inherent variability, we utilized a repeated measure analysis on individual lenses to determine the effects of strain. We determined that the application of 10% axial strain significantly decreases capsule thickness, with thickness reduced on average by 9.9±3.0% (Figure 4E). Following removal of load, capsule thickness returns to unstrained levels (Figure 4D and E).

Increasing axial strains beyond 10% further decreased capsule thickness (Figure S2). The decrease in capsule thickness at 29% axial strain (the highest strain used in this study) is significantly greater than the decrease in capsule thickness at 10% axial strain (p < 0.001). On average, 10% axial strain decreased capsule thickness by 25.8±4.3%. Furthermore, even at 29% axial strain, capsule thickness recovers completely following release from load such that there are no significant difference (p = 0.99) in capsule thickness between unloaded capsules and those recovered from 10% axial strain. These findings extend previous studies on isolated capsules (Fisher, 1969b, a; Krag *et al*., 1997) by demonstrating that capsules are also highly elastic in native, intact, live lens tissues.

### Lens flattening caused by strain leads to a reversible increase in epithelial cell area

Next, we sought to determine if axial strain propagated into the cellular regions of the lens by examining anterior epithelial cells located beneath and attached to the lens capsule. For this examination, we identified a region of interest (ROI) of an identical population of epithelial cells, from images taken prior to, during, and after release from 10% strain (Figure 5A). Measurement of the ROI areas prior to and during strain revealed that the application of 10% axial strain leads to an increase in ROI area (Figure 5A, B).

**Figure 5.**
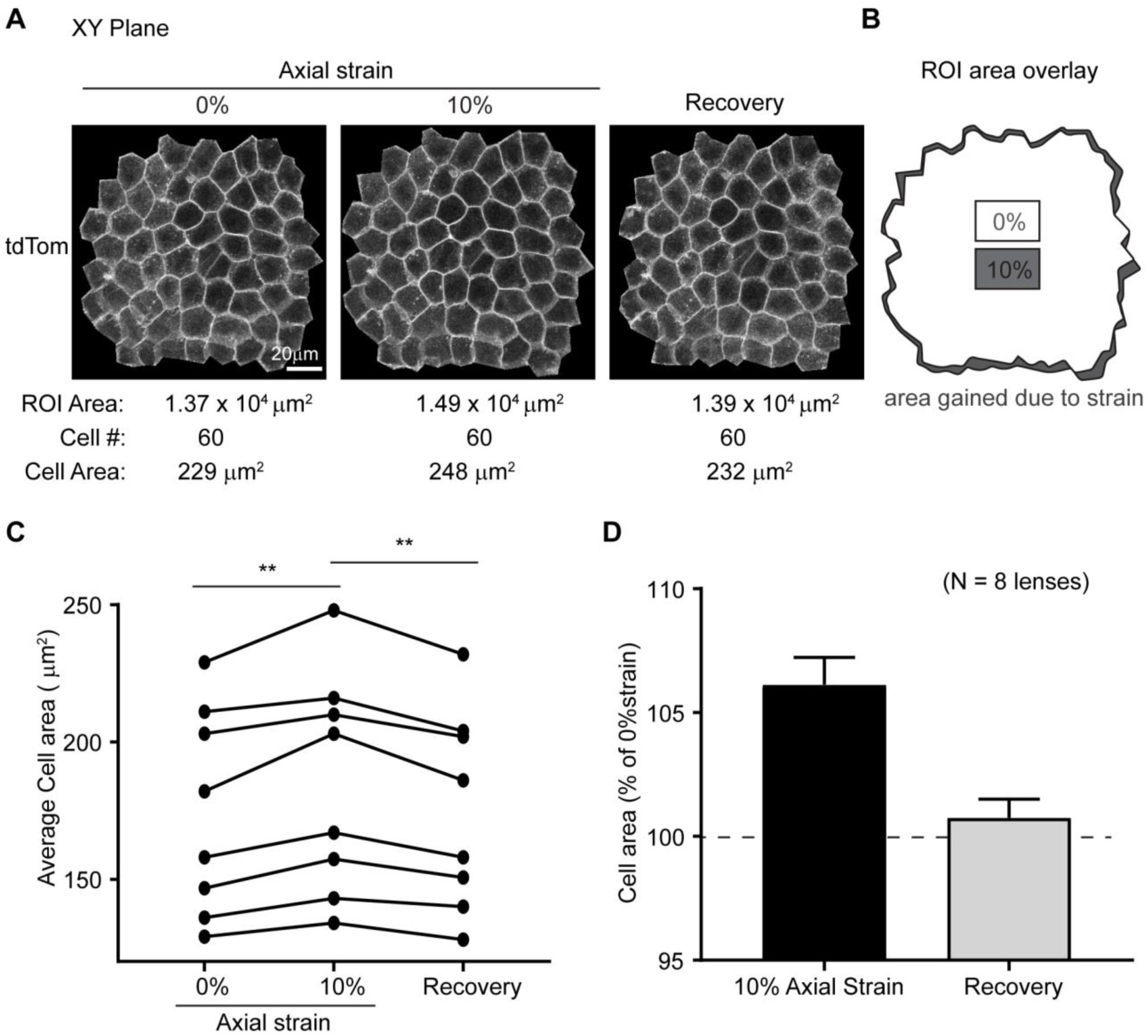
Lens axial strain leads to a reversible increase in anterior epithelial cell area. (A) XY extended view of the mid-region of tdTomato epithelial cells from unstrained (0%), 10% strained, and recovered lenses after release from strain. Total cell area (ROI area) was measured and the average cell area was calculated by dividing ROI area by total cell number as indicated below images. (B) Overlay of cell area at 0% strain (white area) on top of cell area at 10% strain (grey area), demonstrating an increase in area with strain. (C) Repeated measures dot-plots of average epithelial cell area from individual unstrained (0%), 10% strained, and recovered lenses. (F) Average change in epithelial cell area was calculated from data shown in D, normalized to the epithelial area at 0% strain. *, p < 0.05; **, p < 0.01; ***, p < 0.001.

To determine average cell area for individual epithelial cells, the ROI area was divided by the total number of cells within the ROI (Figure 5A). In keeping with a previous study that utilized C57BL/6 wild-type mice of a similar age (Shi *et al*., 2014), we determined that average epithelial cell area is variable between lenses (Figure 5C). Due to this inherent variability in epithelial cell area, a repeated measures analysis on individual lenses was utilized to determine the effects of strain. The application of 10% axial strain significantly increases cell area (Figure 5C) with an average increase in area of 7.0±1.1% (Figure 5D). Furthermore, following removal of 10% axial strain, epithelial cell area returns to unstrained levels (Figure 5C, D). Unlike the capsule where additional strain results in progressive reductions to capsule thickness (Figure S2), there are no further increases in epithelial cell area by increasing strain beyond 10% (Figure S3). The increase in epithelial cell area even at 29% axial strain was not different from the increase in epithelial cell area at 10% axial strain (p = 0.24). Our results demonstrate that axial compression propagates into the epithelial cell layer and epithelial cells expand their areas to near maximal levels at ~10% axial strain.

### Fiber cell tip separation at the suture progressively increases with applied strain and is not completely reversible at high loads

Underlying the anterior epithelial cells, the apical tips of fiber cells meet to form the suture, which is typically a three-branched, Y-shaped suture in mouse lenses (Kuszak *et al*., 2006) (as depicted in Figure 6A). To determine whether the application of axial strain affects the anterior fiber cell tips at the suture, we examined the suture in lenses prior to, during, and after the release from strain. Fiber cell tip separation was quantified by manually tracing the area between fiber cell tips in 3D reconstructions of confocal Z-stacks of lens sutures (Figure S4). We defined the area of this outlined region as the suture gap area.

**Figure 6.**
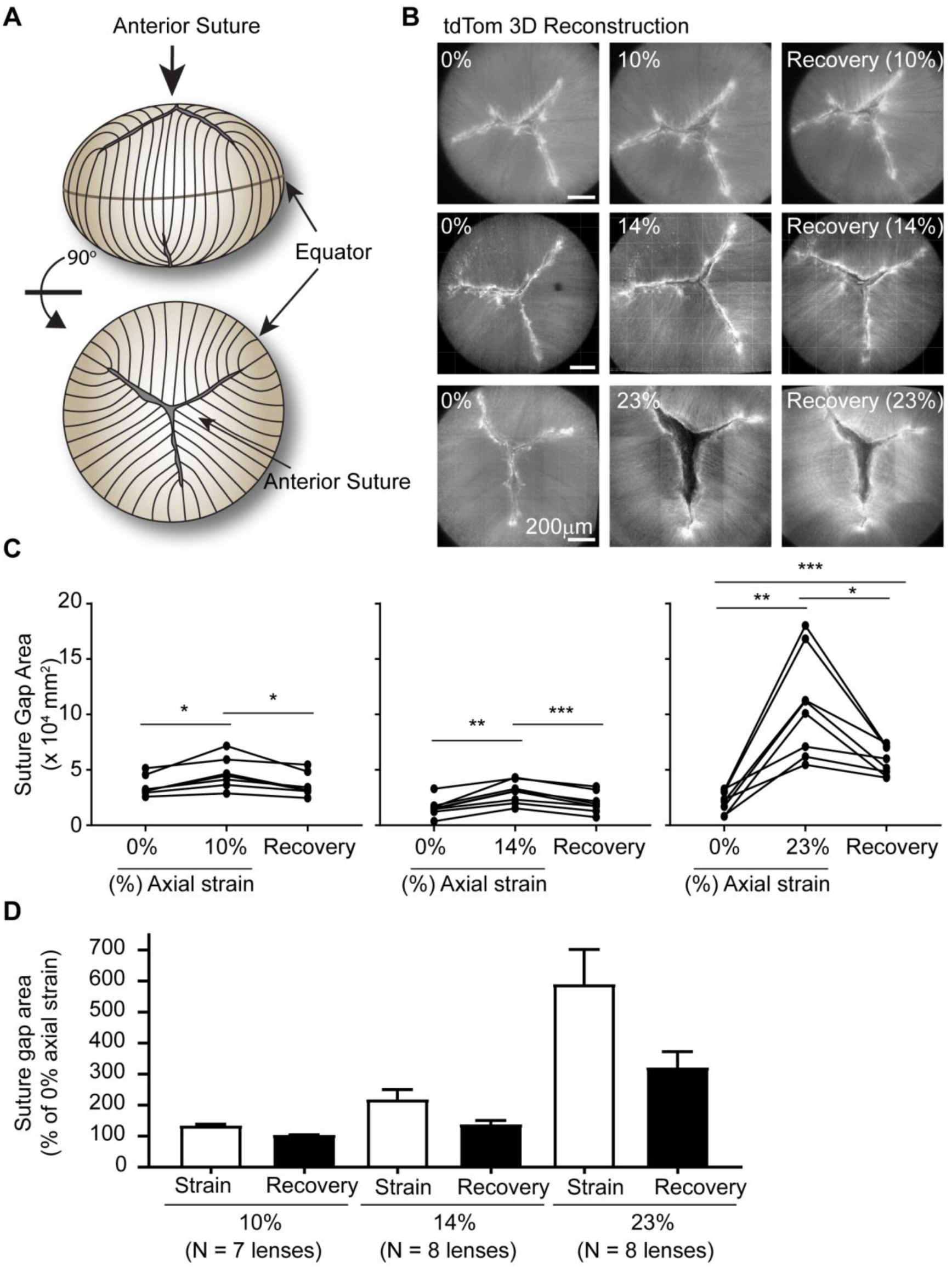
Lens axial strain leads to an irreversible increase in suture gap area at higher strains. (A) Diagram of lens indicating anterior suture in side view and rotated 90° forward for top view of Y-shaped suture. (B) *En face* images of tdTomato (tdTom) fiber cells at anterior suture from 3D reconstructions of confocal Z-stacks in lenses prior to (left column), during (middle column), and after release from (right column) strain. Axial strains of 10% (top row), 14% (middle row) or 23% (bottom row) are shown. The area between opposing fiber cell tips was outlined on images and was defined as the suture gap area (see Figure S4). (C) Repeated measures dot-plots showing the effect of coverslip loading and removal on lens suture gap area for individual lenses. (D) Average change in suture gap area was calculated from data shown in C, normalizing to suture gap area at 0% strain. *, p < 0.05; **, p < 0.01; ***, p < 0.001

We determined that the sutures of tdTomato mouse lenses are variable. Approximately 27% of tdTomato lenses have sutures with four, rather than the typical three, branches (Figure S4). Additionally, the suture gap area is variable between lenses. Therefore, due to the inherent suture variability, a repeated measures analysis on individual lenses was utilized to determine the effects of strain. The application of 10% axial strain significantly increases suture gap area.

Following removal of 10% axial strain, the suture gap area returns to its original size (Figure 6B, C). The amount of fiber cell tip separation at the suture depends on the applied strain (Figure 6D), with axial strains of 10%, 14%, and 23%, the suture gap area increased by 1.4-, 2.2-, and 5.9-fold over the unloaded control, respectively. The suture gap area completely recovers back to its original size after release from 10% or 14% axial strain. However, suture gap area only partially recovers after release from 23% axial strain, with the suture gap remaining 3.2-fold larger than the unstrained control (Figure 6B-D). The application of 29% axial strain also increases suture gap area, which only partially recovers after removal of strain (data not shown). These findings indicate that axial strain affects suture structure in mouse lenses causing separation of fiber tips leading to an increase in the suture gap area. While these effects are reversible at low axial strains (≤14%), they are irreversible at higher axial strains (≥23%), suggesting damage occurs to the suture at higher strains, consistent with previous qualitative observations (Baradia *et al*., 2010).

### Lens flattening caused by strain leads to a reversible increase in equatorial fiber cell width

As lenses are compressed axially, they expand equatorially (Baradia *et al*., 2010; Gokhin *et al*., 2012; Cheng *et al*., 2016a). We determined that at axial strains of 10, 14, 23, and 29% correspond to equatorial strains of 1, 2, 6, and 9%, respectively (Figure 2). At the lens equator, long thin fiber cells are aligned perpendicular to the direction of equatorial strain, therefore fiber cell widths might be expected to expand during axial lens flattening. To examine fiber cell widths, we fixed lenses as they were strained by application of coverslip loads. When fixed under strain, lenses maintain their flattened shape (Figure S5). The utilization of fixed lenses enables orientation of lenses so that the equatorial region is in proximity to the microscope objective, which is necessary for optimum imaging of the individual fiber cells. Fixed lenses were then permeabilized and stained with WGA for the capsule, Hoechst for cell nuclei, and rhodamine-phalloidin for visualization of F-actin at fiber cell membranes (Cheng *et al*., 2017b). The staining of fiber cell membrane-associated F-actin with phalloidin resulted in superior signal-to-noise ratio as compared to the images of fixed tdTomato-labelled fiber cells at the equator (data not shown). This superior labeling of F-actin at membranes is the reason for conducting this portion of the study in wild-type lenses rather than tdTomato lenses.

Since fiber cell widths change with depth in the lens, we standardized the depth at which we measured fiber cell widths in unstrained and strained lenses. For depth standardization, we located the lens fulcrum or modiolus where the apical tips of elongating fiber cells anchor prior to cell elongation (Zampighi *et al*., 2000; Sugiyama *et al*., 2009; Cheng *et al*., 2013), and measured fiber widths within the fiber cell mass at ~10μm inwards from the fulcrum (~4 cell layers deep), as depicted in Figures 7A and 7B. The fulcrum was identified at the equator based on concentrated F-actin staining (Cheng *et al*., 2013) in XZ plane views of sagittal optical sections of 3D reconstructions from confocal Z stacks (red arrow; Figure 7A, C). In single optical sections *en face* (XY plane view), the fulcrum could also be identified by a change in cell organization (revealed by the F-actin at cell boundaries) and by nuclear alignment (red arrow in top panel; Figure 7B, D). Fiber cell widths were measured through line scan analysis of F-actin signal intensity across a line perpendicular to the fiber cell lengths, at the level of the fulcrum, and 10μm deep from the fulcrum. On the resultant line scan profiles of F-actin fluorescence intensity, the periodic signals mark the cell boundaries. These signals were analyzed computationally to obtain high spatial accuracy in determination of fiber widths by peak-to-peak analysis (Gokhin and Fowler, 2017).

**Figure 7.**
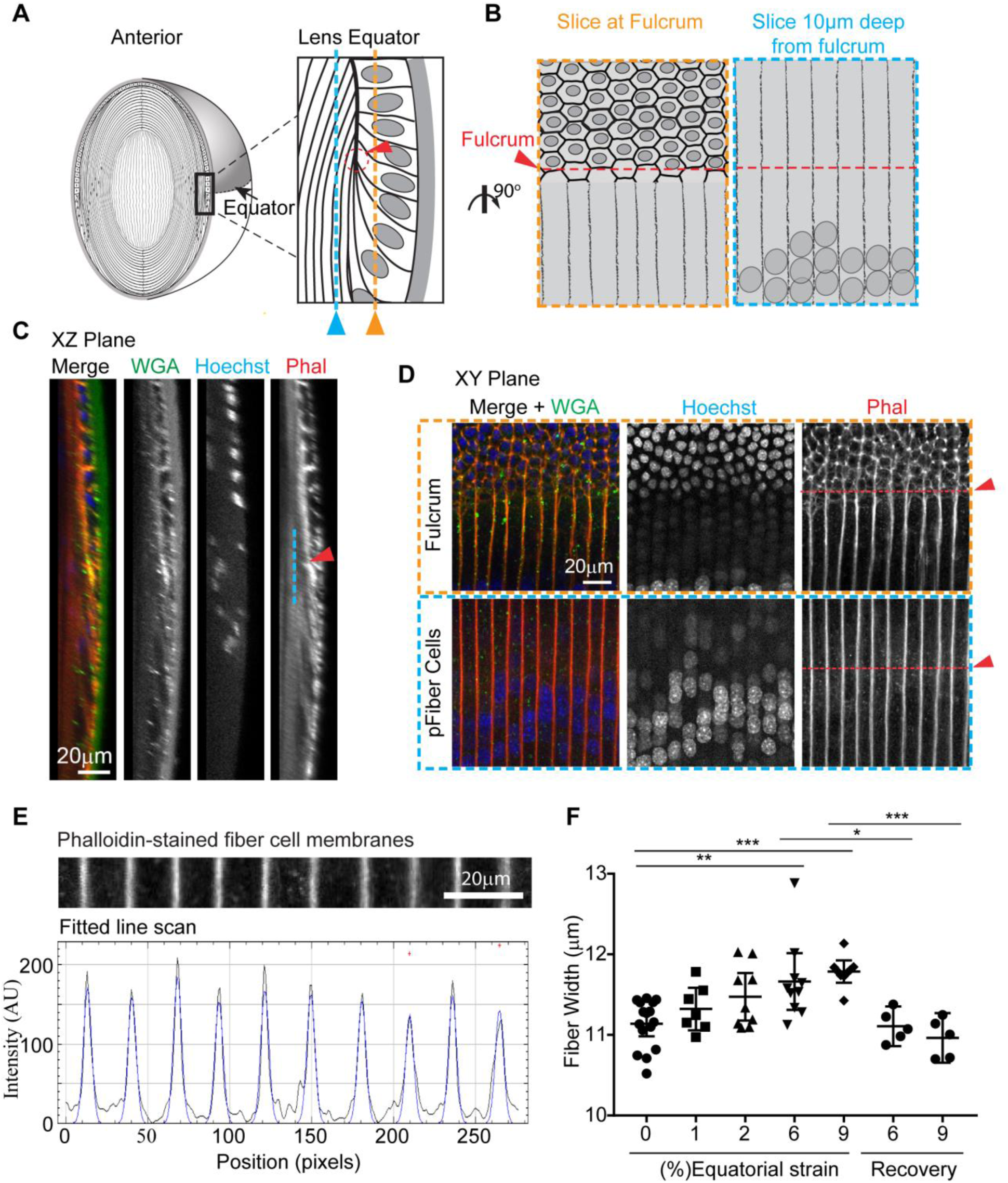
Lens equatorial strains of 6% or 9% lead to increased equatorial fiber widths. (A) Diagram depicting equatorial region of lens (left panel), with equatorial region boxed and magnified (right panel) to indicate the location of the optical section at the epithelial to fiber cell transition (orange) at the fulcrum (red), and the location of the optical section used to measure fiber cell widths 10μm deep from fulcrum (blue). (B) Diagram depicting XY plane of an optical section showing epithelial to fiber transition (left panel outlined in orange dashed box) The fulcrum can be determined by the change in nuclear organization and cell morphology (red arrow and dashed line) and fiber widths were measured at the fulcrum position in an optical section ~10μm deep from fulcrum (right panel outlined in blue dashed box). (C) 3D sagittal reconstruction of confocal Z-stacks, in YZ plane view, of lens equator showing fulcrum (red arrowhead), and peripheral fiber cells ~10μm deep from the fulcrum (blue line). (D) Single optical section (XY plane view) shows the epithelial to fiber transition (top panels) and the peripheral fiber cell layer (pFiber cells) ~10μm deep in this region (bottom panels), as depicted schematically in B. The pFiber cells shown in the bottom panel are from the region marked by blue dashed line in A and B; arrowhead indicates the XZ position of fulcrum. (E) To calculate fiber cell width, fitted-line scan analysis of F-actin intensity across fiber cell membranes at the level of the fulcrum was performed. The distances between F-actin peaks represent fiber cell widths in pixels, which were converted to distance in microns. (F) Dot plots demonstrating the effect of compression on lens fiber width. Each dot represents the average fiber cell width from 1 lens. *, p < 0.05; **, p < 0.01; ***, p < 0.001 as compared to unstrained, 0% strain lenses. Bars, 20 μm (C, D); 10 μm (E).

We determined that, in unstrained lenses, fiber cells are 11.1±0.1μm wide. At 6% equatorial strain, fiber width increased to 11.7±0.2μm (Figure 7F). At 9% equatorial strain, fiber width increased to 11.8±0.1μm. Thus, fiber cell width increased ~4.7% and 5.8% on average, at 6% and 9% equatorial strains on the lens, respectively. Despite trends toward increasing fiber width even at equatorial strains of 1% (p>0.99) or 2% (p=0.15), these were not statistically significant. Following removal of either 6 or 9% equatorial strain, fiber width returned to unstrained levels. Thus, fiber cells at the equator respond to whole lens axial compression by increasing in width, and upon release of strain, fiber cells recover completely.

## Discussion

This study provides new insights into microscale transfer of loads. The distribution of strain differs within the lens as compared to connective soft tissue (tendon, meniscus, annulus fibrosis). In connective tissues, mechanical stress is shielded by the matrix, and strain is largely/completely attenuated prior to reaching the cells (Bruehlmann *et al*., 2004; Fang and Lake, 2015). However, in the eye lens, strain was transferred onto the cells, thereby increasing epithelial cell areas and fiber widths (Figure 8). Lens cells exist as a mass of highly organized and interconnected cells that allows study of force transmission between neighboring cell types and across the bulk tissue. Our work also indicates that microstructures within a tissue respond to mechanical unloading. At low levels of strain (≤14% axial strain), there is complete recovery in bulk lens tissue and all microstructural dimensions. However, at high loads (≥23% axial strain), where bulk tissue recovery is incomplete, there is also incomplete recovery of the suture gap area. The recovery of other microstructures from high loads was not affected. Thus, specific microstructures have different propensities to bear and recover from load.

**Figure 8.**
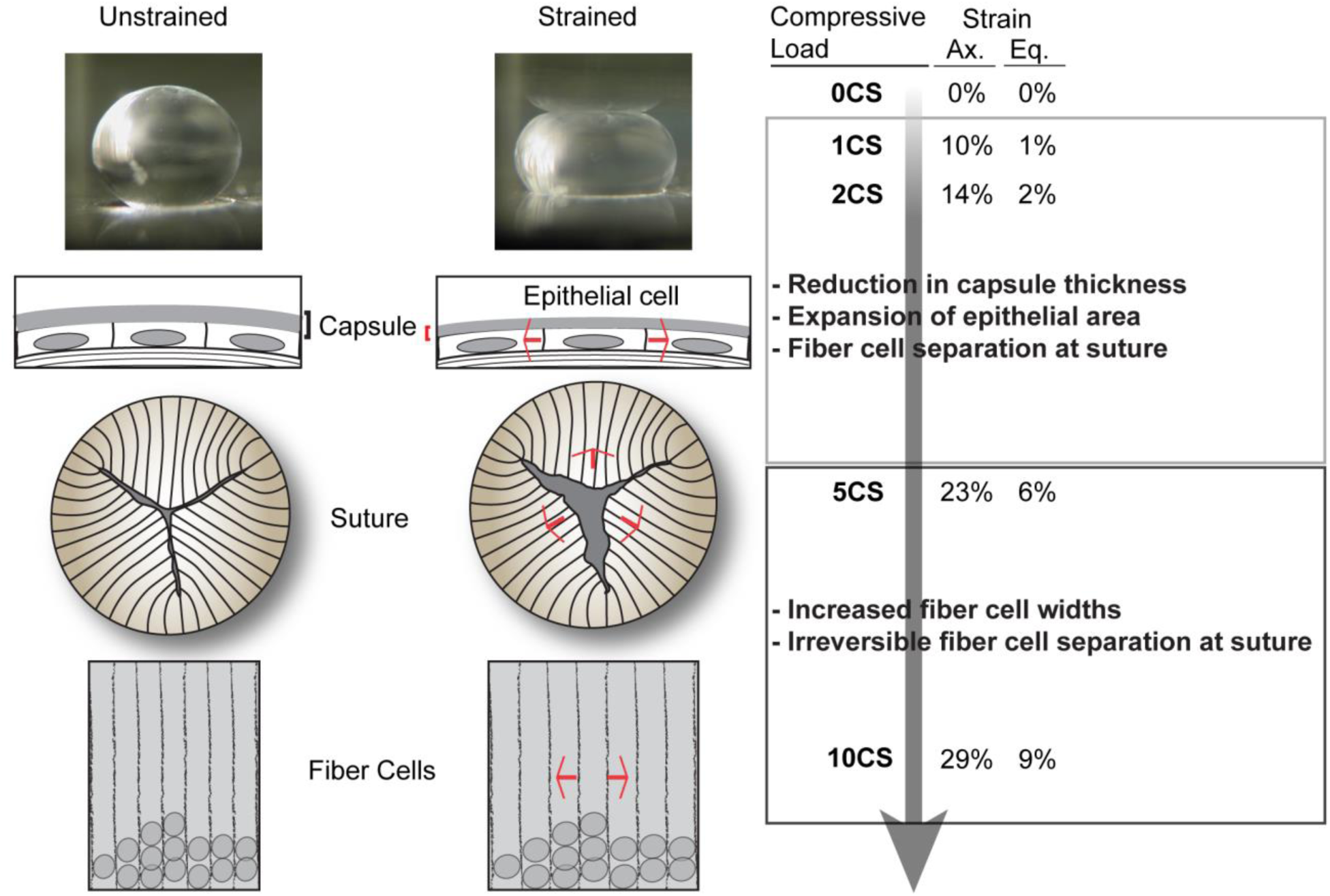
Summary diagram of the effects of axial and equatorial strains on mouse lens shape and capsule and cellular morphologies. Low compressive strains reversibly reduce capsule thickness, increase anterior epithelial cell area, and separation of fiber cell tips at the anterior suture. High compressive strains reversibly increase fiber cell widths at the equator, but irreversibly separated fiber cell tips at the anterior suture.

We find that lens capsule thickness decreases with axial strain and recovers completely following release from strain (Figure 8). The ability of the capsule to reversibly deform with applied strain may not come as a surprise since it has been well documented that isolated capsules are viscoelastic (Fisher, 1969b; Halfter *et al*., 2013). Our observations of intact, and not isolated, lens capsules provide additional support for capsule deformability *in situ*. We also show that capsule thickness decreases progressively with increasing strains. Even at 29% axial strain, the highest level of strain applied, the capsule completely recovers to its original thickness following the release from load. This demonstrates the high propensity for the mouse lens capsule to undergo strain. Intriguingly, in a previous study, it was determined that isolated bovine capsules can undergo up to 76% strain and young human capsules can undergo up to 108% strain (Halfter *et al*., 2013). The mechanical strength and viscoelasticity of lens capsules is thought to be due to type IV collagen polymer structure, a hexagonal network that forms the main component of capsules across species, including mouse and humans (Schmut, 1978; Danysh and Duncan, 2009; Halfter *et al*., 2013).

Loads applied to the anterior of the lens propagate through the lens capsule to the underlying epithelial cells, which respond to 10% bulk axial strain by expanding their cell area (Figure 8). Lens anterior epithelial cells have a specialized network of actin and myosin organized in polygonal arrays which are thought to generate tensional forces to allow for load bearing (Rafferty and Scholz, 1985, 1989; Rafferty *et al*., 1990). Our findings that the epithelial cell area increases with lens flattening is consistent with the idea that the lens epithelial cells are load bearing. Additionally, when lenses are released from 10% strain, the epithelial cell area decreases and returns to its pre-strain area. An interesting question that arises from this data is whether the lens rounding is a consequence of the tension generated by the actin-myosin polygonal arrays in the epithelial cells, as previously hypothesized (Rafferty and Scholz, 1985, 1989; Rafferty *et al*., 1990). Alternatively, the decrease in epithelial cell area may instead be a consequence of forces generated from bulk lens rounding. Although it has been shown that actomyosin disruption can affect the biomechanical response of chick lenses to bear load (Won *et al*., 2015), future experiments that specifically perturb the actin-myosin network in either epithelial or fiber cells of lenses under mechanical stress will be required to address this question. Beyond 10% strain, we did not detect a further increase in epithelial area, possibly due to the variability of anterior epithelial cell areas at higher loads. This variability may arise due to localized epithelial cells sinking into the suture gap at high loads, leading to a decrease in cell area of epithelial cells directly above the suture, but an increase in cell area of epithelial cells away from the suture (data not shown).

Application of load at the lens anterior, resulting in whole lens flattening, causes an increase in fiber cell width at the equator. Notably, unlike the lens capsule and epithelial cells which change their dimensions at low load (1 coverslip), increases in fiber cell width are only detected at higher loads (≥5 coverslips). This could be due to the magnitude of strain being greater axially than equatorially at any given load, as shown here and previously (Cheng *et al*., 2016a). With 1 coverslip, the equatorial strain is only 1 %, whereas axial strain is 10%. Although there is a trend toward small increases in fiber cell widths at 1 and 2% equatorial strain, a power analysis reveals that detection of such small differences would require a very large lens sample size (N = 104 and N = 58 for 1% and 2% equatorial strain, respectively). Nevertheless, we detected an increase in fiber cell widths at 6% and 9% equatorial strain with our lens sample sizes (N = 9 and N= 10 for equatorial strains of 6 and 9%, respectively). The increase in fiber widths may not come as a surprise. The hexagonal shapes and formation of complex interlocking protrusions of the mature fiber cells (Dickson and Crock, 1972; Willekens and Vrensen, 1981; Cheng *et al*., 2016b) imply a cellular architecture optimized for load bearing, by analogy to other hexagonally shaped structures, such as in lung alveoli and trabecular bone, that can undergo large deformations (Gibson *et al*., 1982). We also determined that the increase in fiber cell width is completely reversible following release from strain. The complete recovery of fiber width is consistent with bulk lens equatorial recovery, which was also complete at 6%, and nearly complete at 9% equatorial strain. Since 9% equatorial strain is already beyond the physiological equatorial strain that accommodative lenses would experience *in vivo* (Jackson, 1907; Ziebarth *et al*., 2007; Marussich *et al*., 2015), we did not examine the effect higher loads to determine whether greater equatorial strains led to incomplete recovery.

Fiber cell tip separation at the suture is irreversible at axial strains ≥ 23%. These findings are similar to those of Baradia *et al*. (2010) who showed that in 4-week old C57Bl/6 wild-type mouse lenses, fiber cell tips irreversibly separated when lenses were compressed by 200μm with a specialized confocal microscope objective (Baradia *et al*., 2010). Damage at the suture could explain the incomplete bulk recovery of whole lens shape observed at high strains. These observations of damage raise the question: are mouse lenses, and specifically the fiber cell tips at the sutures, at the ages used in the present study and by Baradia *et al*, still undergoing maturation and therefore unable to tolerate strain? In support of this idea, older more mature fiber cells that are approximately >70μm deep from the anterior surface do not appear to separate at the suture (data not shown). However, it is unclear if this is due solely to fiber cell maturity or to dissipation of loads with distance inwards. Therefore, examination of lenses from older mice may give more direct insight into this question.

This study provides a framework to examine multiscale transfer of strain in lenses of other species. Mouse and primate lenses have similar biomechanical characteristics, including the increase in lens modulus with age (Manns *et al*., 2007; Ziebarth *et al*., 2007; Scarcelli *et al*., 2011; Cheng *et al*., 2016a), as well as similar capsule and cellular structures (Wanko and Gavin, 1959; Rafferty and Scholz, 1985; Kuszak *et al*., 2006; Danysh and Duncan, 2009). Thus, mouse lenses are a suitable starting point to develop methodologies and determine characteristics of multiscale transfer. Mice also can be genetically engineered and the use of tdTomato mice was advantageous to allow for the identification of microstructures in live lenses in this study. However, mouse lenses are presumed to have limited ability for shape change *in vivo* (Rafferty and Scholz, 1989; Baradia *et al*., 2010). There are microstructural differences between mouse and accommodating (primate) lenses (Kuszak *et al*., 2006; Kuszak and Zoltoski, 2006). Whereas mouse lenses typically have sutures with three or four branches, accommodative primate lenses have sutures shaped in a complex star pattern with multiple branches. In addition, mouse lens fiber cells form aligned sutures between concentric layers, while in primate lenses, suture patterns are offset between concentric shells. Therefore, loads may be more effectively transmitted across the multiple suture branches and offset layers in primate lenses. Furthermore, scanning electron microscopy studies show that lens fiber cell tip morphologies in primate lenses and mouse lenses are different. The fiber cell tips in accommodative lenses taper at their ends and have been proposed to allow fiber cell tips to slide over one another during lens shape changes (Kuszak and Zoltoski, 2006). Mouse lens fiber cell tips, on the other hand, have little taper and limited overlap and are, therefore, unlikely to slide over each other (Kuszak *et al*., 2006; Kuszak and Zoltoski, 2006). The lower number of suture branches and reduced tip adhesive area in mouse lens fiber cell tips could explain why the mouse lens suture separates at high loads. Studies evaluating the effect of load on suture morphology and fiber cell tips in primate lenses, and whether any damage occurs to the suture under load, will provide insight into species-specific differences.

In conclusion, our study establishes a relationship between whole tissue biomechanics and specific microstructures in the lens model. Gross lens shape change caused by strain results microstructural level changes in all the regions of the lens that we examined. This provides critical information that will be useful in expanding current biomechanical models of lens shape changes (Reilly and Ravi, 2010; Reilly, 2014; Wang *et al*., 2017). Furthermore, the identification of a relationship between microstructure and tissue biomechanics supports the idea that age-dependent microstructural alterations, such as an increase in stiffness of cells or matrix (Starodubtseva, 2011; Lacolley *et al*., 2017), could have biomechanical consequences at the tissue level. Our methodology could be adapted to further determine which specific molecules, present in either matrix or cells, affect soft tissue biomechanics. Thus, the lens is a novel robust model that will allow in-depth studies to elucidate the complexities of soft tissue multiscale load transfer.

## Materials and Methods

### Mice

Wild-type and Rosa26-tdTomato mice tandem dimer-Tomato (B6.129(Cg)-Gt(ROSA) (tdTomato) (Jackson Laboratories) in the C57BL/6J background between the ages of 6-9 weeks were used for experiments. The tdTomato mice express tdTomato protein fused in frame to connexin 43 (Muzumdar *et al*., 2007). Genotyping confirmed that the mice were wild-type for Bfsp2/CP49. Breeding and care of mice were conducted in accordance with an approved animal protocol from The Scripps Research Institute. Following euthanization, eyes were enucleated from mice and placed in a dissection dish in phosphate buffered saline (PBS) (137mM NaCl, 2.7mM KCl, 8.1mM Na_2_HPO4, 1.5mM KH_2_PO4; pH 8.1). All experiments were performed at room temperature in the above mentioned PBS. We did not observe any obvious changes in lens transparency during experiments.

### Lens dissections

To investigate bulk mechanical properties of lenses as well as to image the equatorial regions of fixed lenses, complete lens dissection was performed (Cheng *et al*., 2016a). Lenses were dissected from mouse eyes under oblique illumination of a dissecting microscope. Briefly, the optic nerve was removed from freshly enucleated eyes using microdissection scissors. Then, fine straight tweezers were inserted into the remaining hole where the optic nerve entered the eye and scissors were used to cut from the posterior half way around the eye to the anterior region. Finally, the lens was then gently pushed out by applying pressure to the uncut side of the eye.

To investigate the anterior regions of the live lens, partial lens dissection was performed. Partial lens dissection minimizes mechanical damage that can occur at the anterior region with complete dissection. For partial lens dissection, an incision was first made at the cornea-sclera junction of the eye using the tip of a scalpel blade (#11, Feather Safety Razor Co, Osaka, Japan). Next, microdissection scissors was inserted into the incision, and small cuts were made all the way around the eye along cornea-sclera junction. Following removal of the cornea, the iris was gently peeled off the lens using fine forceps to expose the anterior region (Figure S6; anterior view). Next, the optic nerve was cut off using scissors as close to the eye as possible. Finally, scissors were inserted through the hole where the optic nerve entered the eye and utilized to cut half the way up the posterior region of the eye and then circumferentially around the eye equator exposing the posterior regions of the lens (Figure S6; posterior view). Following this partial lens dissection, the eye lens with an attached band of the surrounding equatorial eye tissue remained (Figure S6; side view).

### Biomechanical compressive testing of lenses

Bulk compressive properties of lenses were assessed by sequential application of glass coverslips onto the completely dissected lens as previously described (Gokhin *et al*., 2012; Cheng *et al*., 2016a). Briefly, lenses were placed in PBS within a divot of a custom-made loading chamber that was situated underneath a dissecting microscope. Side views of the lens were captured through a 45° angled mirror, placed at a fixed distance from lens. Images were acquired through a digital camera mounted onto the microscope. Coverslips, with an average weight of 129.3mg, were applied sequentially, one-at-time, onto the lenses. The lenses were compressed for 2 minutes to allow for stress-relaxation equilibration, prior to image acquisition. The axial and equatorial diameters were measured on images using FIJI software. Lens diameters were converted from pixels to metric (mm) units using the mirror edge (length of 5mm) for calibration. Percentage strain was calculated using the formula ɛ = 100 × (d_1_−d_o_)/d_o_, where ɛ is either axial or equatorial strain, d_l_ is the axial or equatorial diameter at a given load, and d_o_ is the axial or equatorial diameter at the unstrained (0% strain) state. Lens bulk recovery or resilience was calculated from axial diameter values using the formula recovery = d_f_/d_o_ where d_f_ is the axial or equatorial diameter of the lens following the release from a given load.

### Effect of compression on anterior region of lens

Confocal examination of lens anterior regions under compression was performed on live, partially dissected, tdTomato lenses. Immediately following dissection, lenses were stained with fluorescent CF640 dye-conjugated to wheat germ agglutinin (WGA) (Biotium, Fremont, CA) (1:100) and Hoechst 33342 (Biotium) (1:500) in PBS to visualize the lens capsule and epithelial cell nuclei, respectively. After 15 minutes, stained live lenses were then transferred onto glass-bottomed culture dishes (10 mm microwell; MatTek Corp., Ashland, MA) in 3mL of PBS supplemented with 1.8units of Oxyrase (Oxyrase inc., Mansfield, OH). To prevent movement of lenses during imaging, lenses were immobilized in the culture dishes within a 2mm diameter circular divot that was created, using a biopsy punch, in a thin layer of agarose (4%w/v in PBS). Unconfined compression was applied by sequential application of microscope glass coverslips onto the lenses.

### Effect of compression on equatorial region of lens

The effect of strain on equatorial fiber cell widths was investigated in fixed lenses. Completely dissected lenses were placed in 3mL of PBS and immobilized within an agarose divot, as above. Coverslips were then applied to compress the lenses, as above. After 2 minutes of compression, the strained lenses were fixed by adding 1 mL of 16% paraformaldehyde to PBS (to achieve a final concentration of 4% paraformaldehyde). After 30 minutes, coverslips were removed, and lenses were incubated in the 4% paraformaldehyde fixative for an additional 90 minutes. Fixed lenses were washed in PBS, placed in permeabilization/blocking solution for 30 minutes, and then stained with WGA (1:100), rhodamine-phalloidin conjugate (1:20) and Hoechst 33342 (1:500) for 2 hours. Lenses were washed in PBS, three times for 5 minutes, and then immobilized on their equatorial edges by placing within an agarose wedge (Cheng *et al*., 2017a).

### Confocal microscopy and image analysis

Imaging was performed on a Zeiss 880 laser-scanning confocal microscope (Zeiss, Germany) using a 10x air C-Apo W 0.45NA objective, a 20x air Plan Apo 0.8NA objective, or a 40x oil Plan Apo 1.4NA objective. Z-stacks were acquired at a digital zoom of 0.6 or 1.0, with Z-step sizes of 0.7μm (10x or 20x objective) and 0.3μm (40x objective). Raw images were processed using either Zen (Zeiss), or FIJI software. 3D reconstructions were performed on Z-stack data using Arivis Vision4D (Arivis AG) software. Morphometric image analysis was performed on FIJI software. Detailed explanations of the morphometric analysis are provided within the Results section. Briefly, capsule thickness was calculated by line scan analysis of XZ plane view images to obtain intensity distributions of WGA staining (capsule) and tdTomato labeling (cell membranes) from the outside of the lens across the capsule to the epithelium. To measure epithelial cell area, a region of interest (ROI), containing an identical population of cells, was identified in XY images that were taken prior to, during, and after the release from strain. Average cell area was calculated by dividing the ROI area by the total number of cells within the ROI. Suture gap areas were measured by manually outlining sutures on 3D reconstructed images of tdTomato-labeled fiber cells from the outer layers down to 100μm depths. Fiber cell widths of rhodamine phalloidin-stained fiber cell membranes were determined at the lens equator, ~10μm inwards from the fulcrum, using FIJI with Distributed Deconvolution (Ddecon) (available at: http://www.scripps.edu/fowler/) plugin. Ddecon deconvolutes fluorescent line scan profiles into a Gaussian point-spread function using a mathematical algorithm for deconvolution. The mathematical algorithm is based on the diffractive properties of light and predictive models providing high spatial precision (Gokhin and Fowler, 2017). The ‘Z-line’ predictive model was utilized in the analysis of fiber cell width.

### Statistical Analysis

To detect differences between two groups of data, paired T-tests were utilized. Differences between multiple groups in live lenses were detected using a one-way repeated measures analysis of variance followed by a multiple comparison’s test with Sidak’s adjustment, post-hoc. To analyze differences between multiple groups in fixed lenses a one-way analysis of variance followed by a multiple comparison’s test with Bonferroni adjustment was utilized, post-hoc.

## Acknowledgements.

We would like to thank Michael Amadeo, Dorothy DeBiasse, and Alyssa Mendenhall for assistance with assessment of lens biomechanics.

## Supplemental Figure Legends

**Figure S1.**
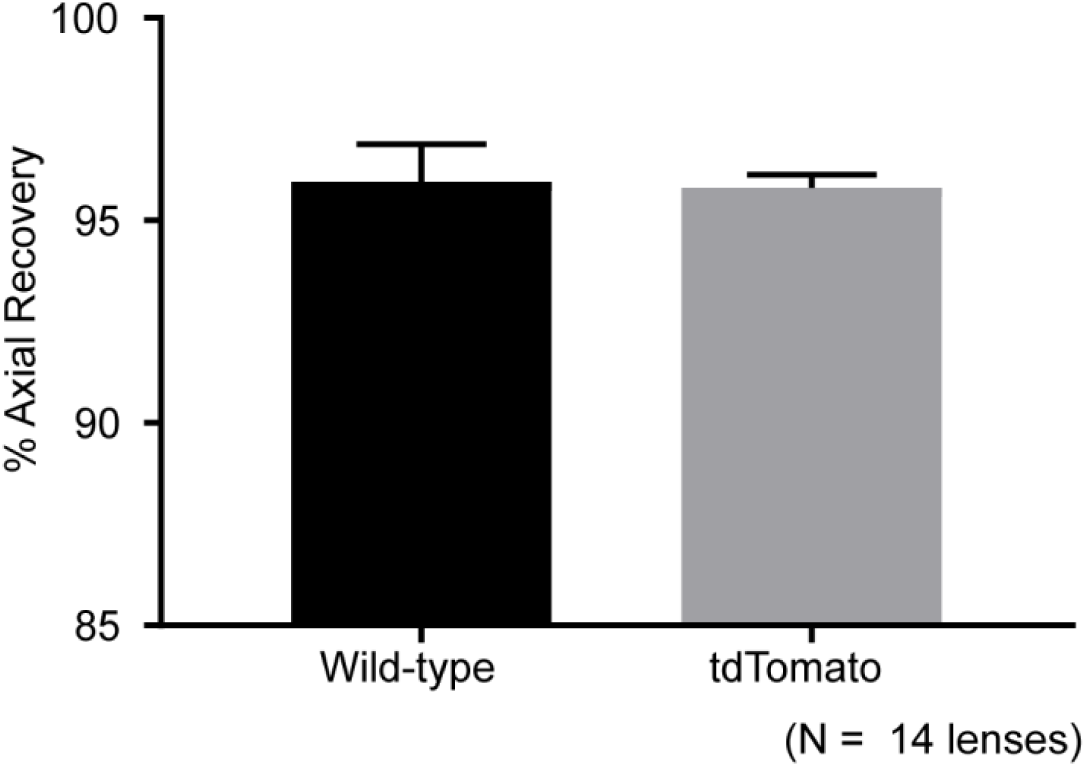
Average axial recovery of wild-type and tdTomato mouse lenses following 29% axial strain. There is no difference in recovery between wild-type and tdTomato lenses. (N = 7 lenses for each genotype)

**Figure S2.**
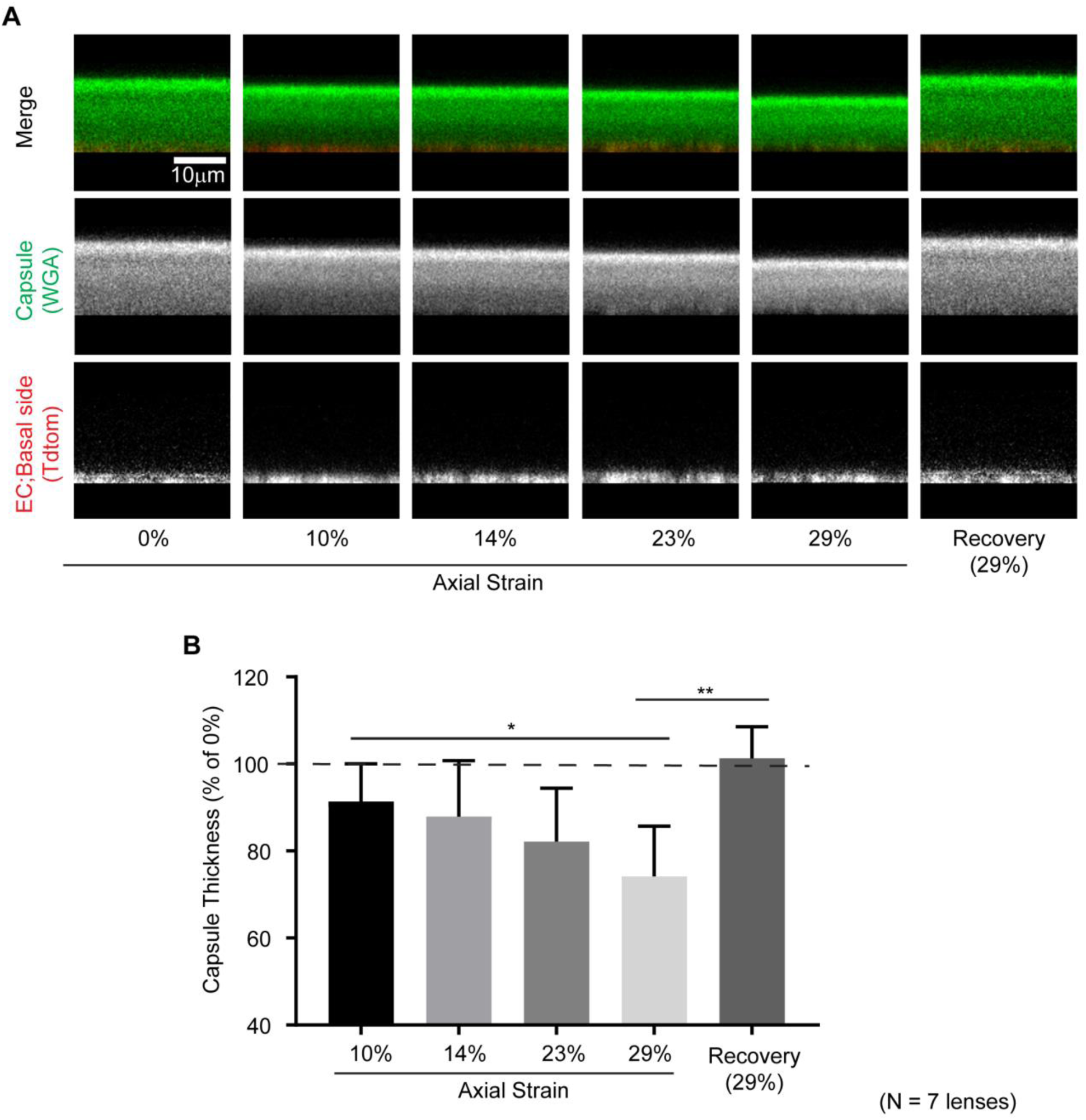
The effect of 10-29% axial strains on capsule thickness for tdTomato lenses. (A) Sagittal (XZ plane view) reconstruction of confocal Z-stacks of tdTomato (red) lenses stained with WGA (green) to visualize capsules. Scale bar, 10μm. (B) Average capsule thickness progressively decreased with increasing axial strain. At the highest axial strain (29%), the capsules recovered completely to unstrained levels. (N = 7)

**Figure S3.**
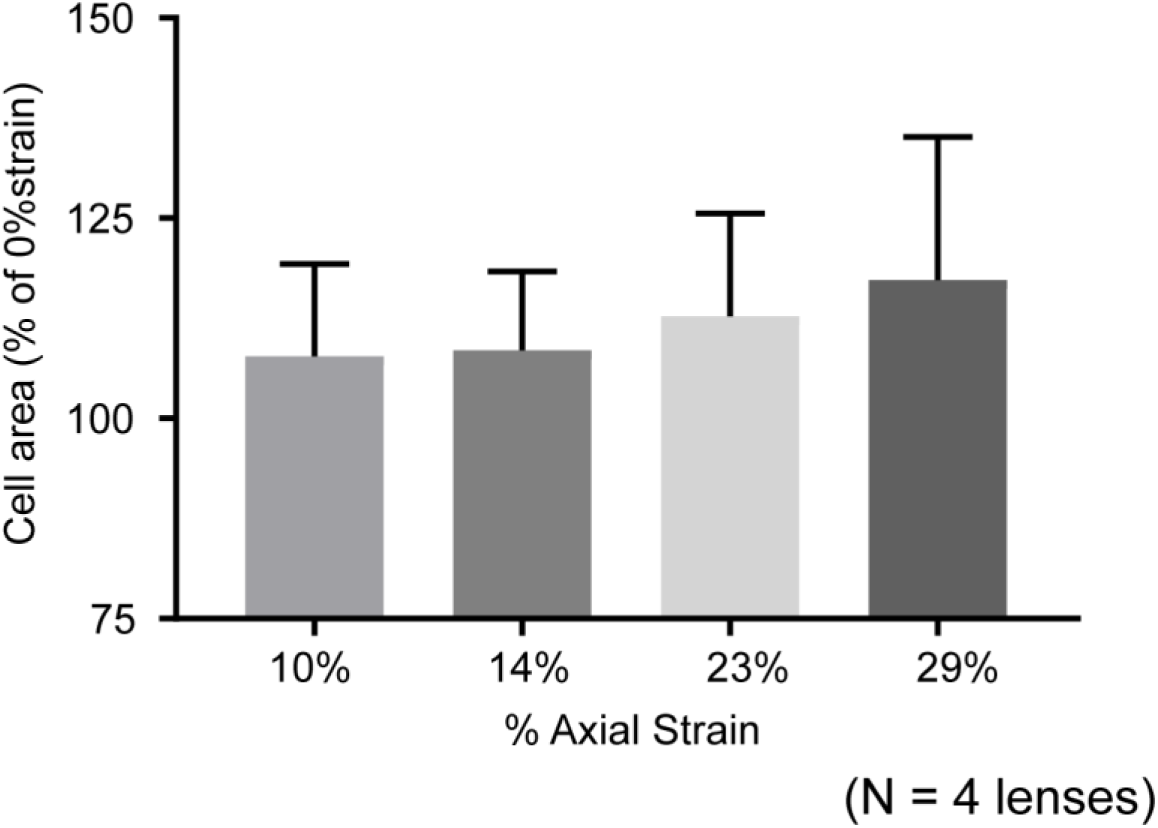
The effect of 10-29% axial strain on epithelial cell area. Average epithelial cell area was similar between 10% axial strain and axial strains greater than 10%. Differences were not statically significant. (N = 4)

**Figure S4.**
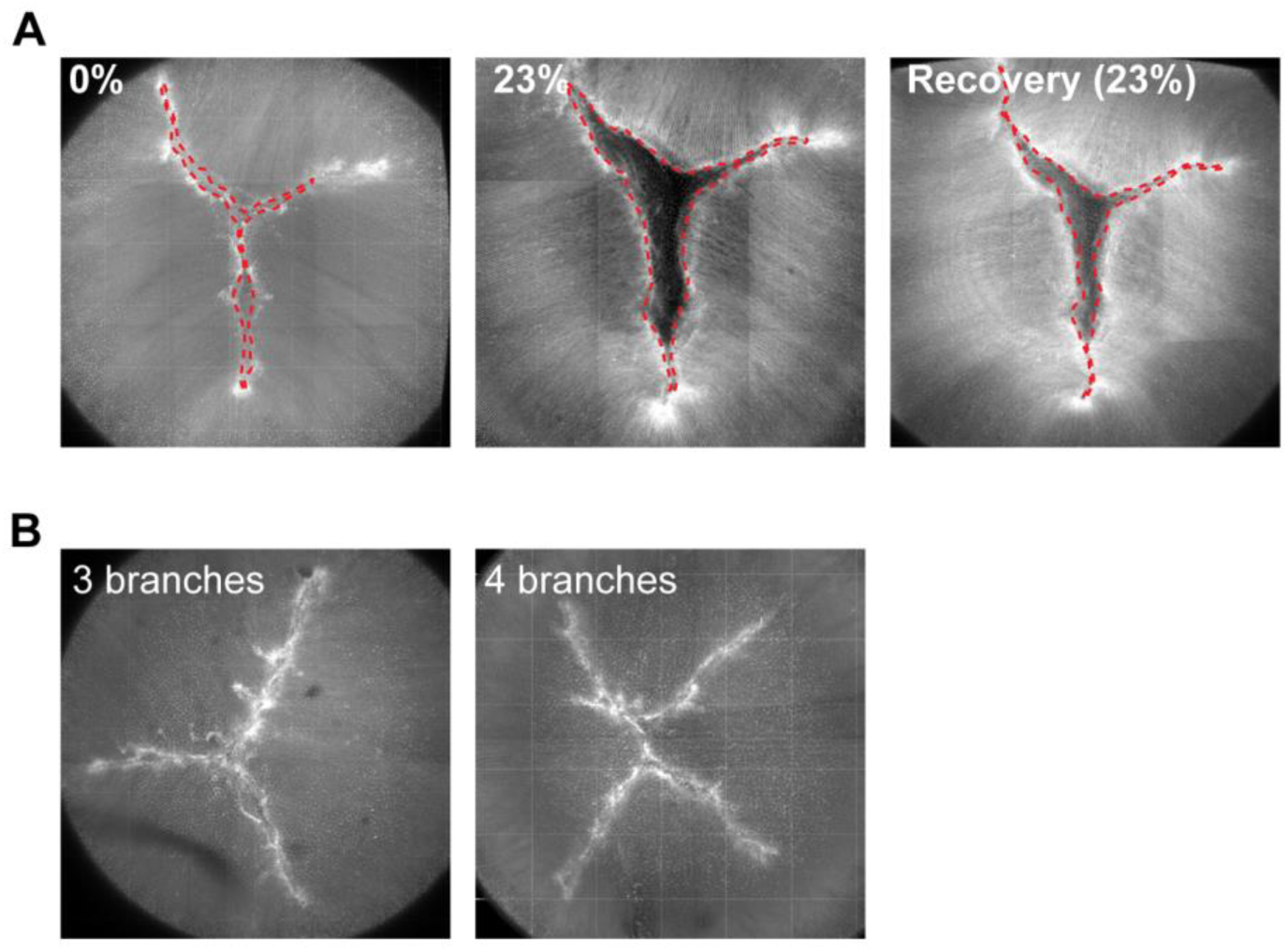
Quantification of fiber cell tip separation through suture gap area analysis. (**A**) Suture gap areaswere measured by manual tracing of fiber tips on *en face* images of anterior suture from 3D reconstructions of confocal Z-stacks in unstrained (left column), strained (middle column), and recovered lenses (right column). (B) Images showing the typical Y-shape three-branched suture seen in ~63% of mouse lenses (left column) and four-branched suture seen in ~27% of tdTomato lenses (right column).

**Figure S5.**
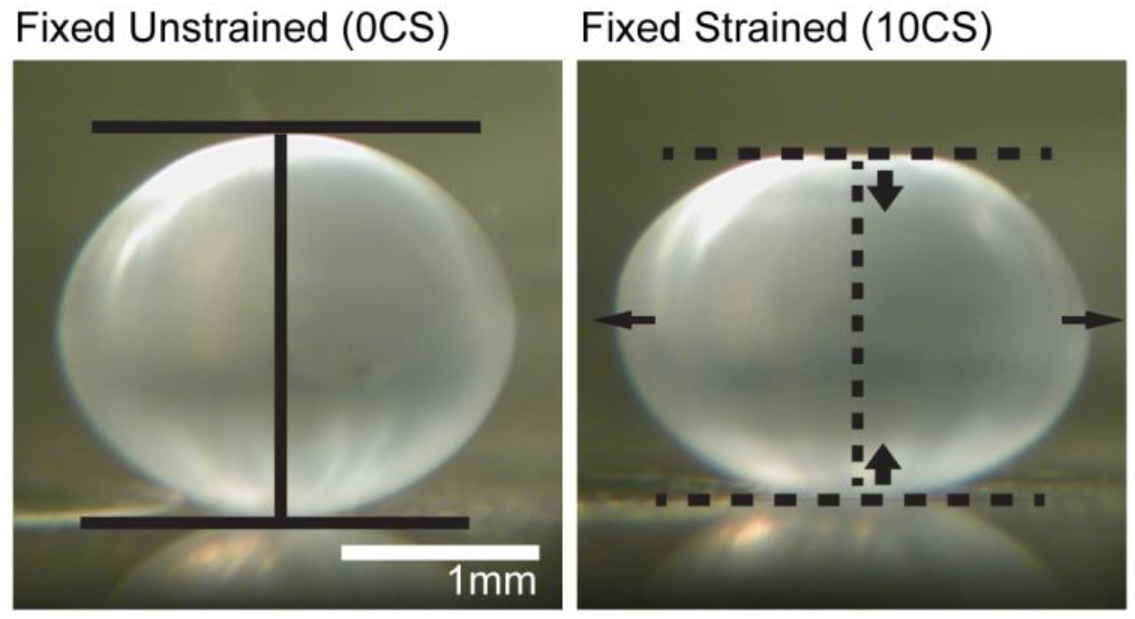
Fixing lenses in the strained state with paraformaldehyde maintains the flattened lens shape. Side view images of wild-type mouse lens fixed in the unstrained (left column) or strained with 10 coverslips (right column). Solid and dashed lines indicate the surfaces of the unstrained and strained lenses, respectively.

**Figure S6.**
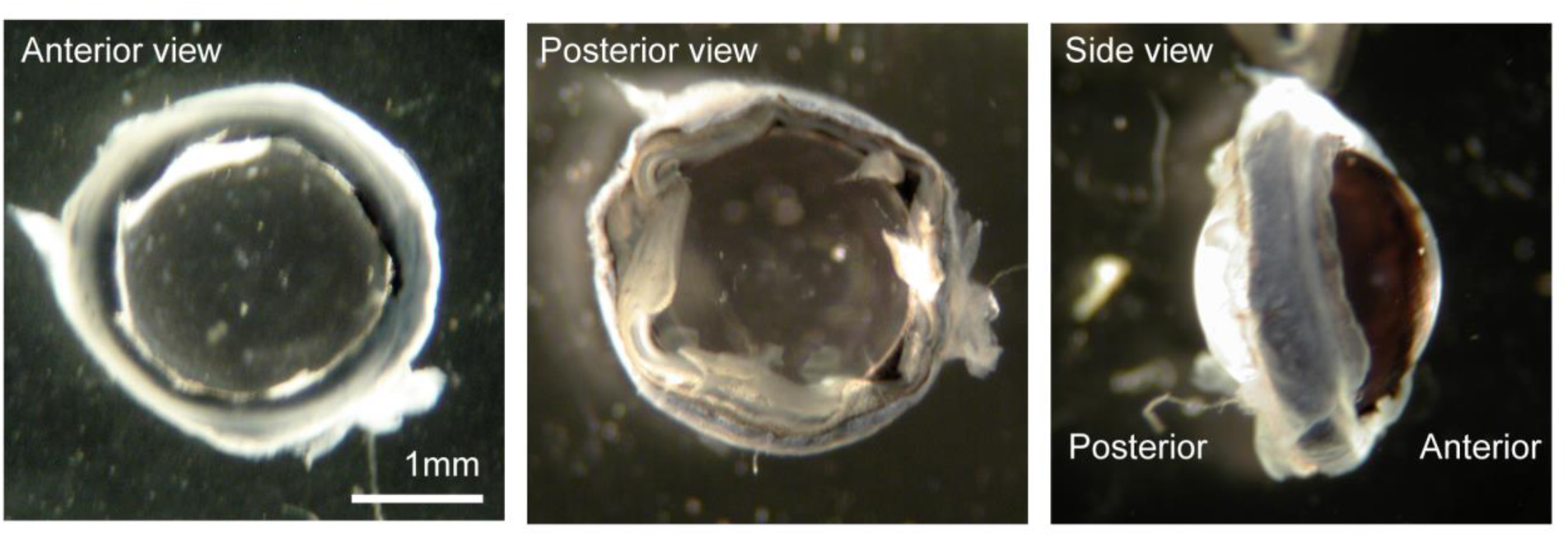
Photos depicting partial dissection of mouse lenses. (A) The anterior region of the lens is exposed by cutting along the cornea-sclera junction. (B) The posterior region of the lens is exposed by cutting to the equator of the eye from the posterior and then around the equator of the eye. (C) Side view showing that partial dissection results in a band of tissue (consisting of sclera, ciliary body and muscle, iris, and zonules) that remains surrounding the lens. Scale bar, 1mm.

## References

Bailey, S.T., Twa, M.D., Gump, J.C., Venkiteshwar, M., Bullimore, M.A., and Sooryakumar, R. (2010). Light-scattering study of the normal human eye lens: elastic properties and age dependence. IEEE Trans Biomed Eng 57, 2910–2917.

Baradia, H., Nikahd, N., and Glasser, A. (2010). Mouse lens stiffness measurements. Exp Eye Res 91, 300–307.

Bonakdar, N., Gerum, R., Kuhn, M., Sporrer, M., Lippert, A., Schneider, W., Aifantis, K.E., and Fabry, B. (2016). Mechanical plasticity of cells. Nat Mater 15, 1090–1094.

Bruehlmann, S.B., Hulme, P.A., and Duncan, N.A. (2004). In situ intercellular mechanics of the bovine outer annulus fibrosus subjected to biaxial strains. J Biomech 37, 223–231.

Campbell, R.E., Tour, O., Palmer, A.E., Steinbach, P.A., Baird, G.S., Zacharias, D.A., and Tsien, R.Y. (2002). A monomeric red fluorescent protein. Proc Natl Acad Sci U S A 99, 7877–7882.

Candiello, J., Cole, G.J., and Halfter, W. (2010). Age-dependent changes in the structure, composition and biophysical properties of a human basement membrane. Matrix Biol 29, 402–410.

Cheng, C., Ansari, M.M., Cooper, J.A., and Gong, X. (2013). EphA2 and Src regulate equatorial cell morphogenesis during lens development. Development 140, 4237–4245.

Cheng, C., Fowler, V.M., and Gong, X. (2017a). EphA2 and ephrin-A5 are not a receptor-ligand pair in the ocular lens. Exp Eye Res 162, 9–17.

Cheng, C., Gokhin, D.S., Nowak, R.B., and Fowler, V.M. (2016a). Sequential Application of Glass Coverslips to Assess the Compressive Stiffness of the Mouse Lens: Strain and Morphometric Analyses. J Vis Exp.

Cheng, C., Nowak, R.B., Biswas, S.K., Lo, W.K., FitzGerald, P.G., and Fowler, V.M. (2016b). Tropomodulin 1 Regulation of Actin Is Required for the Formation of Large Paddle Protrusions Between Mature Lens Fiber Cells. Invest Ophthalmol Vis Sci 57, 4084–4099.

Cheng, C., Nowak, R.B., and Fowler, V.M. (2017b). The lens actin filament cytoskeleton: Diverse structures for complex functions. Exp Eye Res 156, 58–71.

Danysh, B.P., Czymmek, K.J., Olurin, P.T., Sivak, J.G., and Duncan, M.K. (2008). Contributions of mouse genetic background and age on anterior lens capsule thickness. Anat Rec (Hoboken) 291, 1619–1627.

Danysh, B.P., and Duncan, M.K. (2009). The lens capsule. Exp Eye Res 88, 151–164.

Desrochers, J., and Duncan, N.A. (2010). Strain transfer in the annulus fibrosus under applied flexion. J Biomech 43, 2141–2148.

Dickson, D.H., and Crock, G.W. (1972). Interlocking patterns on primate lens fibers. Invest Ophthalmol 11, 809–815.

Doyle, L., Little, J.A., and Saunders, K.J. (2013). Repeatability of OCT lens thickness measures with age and accommodation. Optom Vis Sci 90, 1396–1405.

Dumont, S., and Prakash, M. (2014). Emergent mechanics of biological structures. Mol Biol Cell 25, 3461–3465.

Fang, F., and Lake, S.P. (2015). Multiscale strain analysis of tendon subjected to shear and compression demonstrates strain attenuation, fiber sliding, and reorganization. J Orthop Res 33, 1704–1712.

Fisher, R.F. (1969a). Elastic constants of the human lens capsule. J Physiol 201, 1–19.

Fisher, R.F. (1969b). The significance of the shape of the lens and capsular energy changes in accommodation. J Physiol 201, 21–47.

Gibson, L.J., Ashby, F.R.S., Schajer, G.S., and Robertson, C.I. (1982). The mechanics of two-dimensional cellular materials. Proceedings of the Royal Society of London. A. Mathematical and Physical Sciences 382, 25–42.

Gokhin, D.S., and Fowler, V.M. (2017). Software-based measurement of thin filament lengths: an open-source GUI for Distributed Deconvolution analysis of fluorescence images. J Microsc 265, 11–20.

Gokhin, D.S., Nowak, R.B., Kim, N.E., Arnett, E.E., Chen, A.C., Sah, R.L., Clark, J.I., and Fowler, V.M. (2012). Tmod1 and CP49 synergize to control the fiber cell geometry, transparency, and mechanical stiffness of the mouse lens. PLoS One 7, e48734.

Halfter, W., Candiello, J., Hu, H., Zhang, P., Schreiber, E., and Balasubramani, M. (2013). Protein composition and biomechanical properties of in vivo-derived basement membranes. Cell Adh Migr 7, 64–71.

Han, W.M., Heo, S.J., Driscoll, T.P., Smith, L.J., Mauck, R.L., and Elliott, D.M. (2013). Macro- to microscale strain transfer in fibrous tissues is heterogeneous and tissue-specific. Biophys J 105, 807–817.

Heidemann, S.R., and Wirtz, D. (2004). Towards a regional approach to cell mechanics. Trends Cell Biol 14, 160–166.

Heys, K.R., Cram, S.L., and Truscott, R.J. (2004). Massive increase in the stiffness of the human lens nucleus with age: the basis for presbyopia? Mol Vis 10, 956–963.

Hozic, A., Rico, F., Colom, A., Buzhynskyy, N., and Scheuring, S. (2012). Nanomechanical characterization of the stiffness of eye lens cells: a pilot study. Invest Ophthalmol Vis Sci 53, 2151–2156.

Jackson, E. (1907). Amplitude of Accommodation at Different Periods of Life. Cal State J Med 5, 163–166.

Krag, S., Olsen, T., and Andreassen, T.T. (1997). Biomechanical characteristics of the human anterior lens capsule in relation to age. Invest Ophthalmol Vis Sci 38, 357–363.

Kuszak, J.R., Bertram, B.A., Macsai, M.S., and Rae, J.L. (1984). Sutures of the crystalline lens: a review. Scan Electron Microsc, 1369–1378.

Kuszak, J.R., Mazurkiewicz, M., Jison, L., Madurski, A., Ngando, A., and Zoltoski, R.K. (2006). Quantitative analysis of animal model lens anatomy: accommodative range is related to fiber structure and organization. Vet Ophthalmol 9, 266–280.

Kuszak, J.R., and Zoltoski, R.K. (2006). Focus on Eye Research: The mechanism of accommodation at the fiber level (p.117–132). Nova Science Publishers, Inc.

Lacolley, P., Regnault, V., Segers, P., and Laurent, S. (2017). Vascular Smooth Muscle Cells and Arterial Stiffening: Relevance in Development, Aging, and Disease. Physiol Rev 97, 1555–1617.

Lampi, M.C., and Reinhart-King, C.A. (2018). Targeting extracellular matrix stiffness to attenuate disease: From molecular mechanisms to clinical trials. Sci Transl Med 10.

Lovicu, F.J., and Robinson, M.L. (2004). Development of the ocular lens. Cambridge University Press: Cambridge, UK; New York.

Manns, F., Parel, J.M., Denham, D., Billotte, C., Ziebarth, N., Borja, D., Fernandez, V., Aly, M., Arrieta, E., Ho, A., and Holden, B. (2007). Optomechanical response of human and monkey lenses in a lens stretcher. Invest Ophthalmol Vis Sci 48, 3260–3268.

Marussich, L., Manns, F., Nankivil, D., Maceo Heilman, B., Yao, Y., Arrieta-Quintero, E., Ho, A., Augusteyn, R., and Parel, J.M. (2015). Measurement of Crystalline Lens Volume During Accommodation in a Lens Stretcher. Invest Ophthalmol Vis Sci 56, 4239–4248.

Muzumdar, M.D., Tasic, B., Miyamichi, K., Li, L., and Luo, L. (2007). A global double-fluorescent Cre reporter mouse. Genesis 45, 593–605.

Oxlund, H., and Andreassen, T.T. (1980). The roles of hyaluronic acid, collagen and elastin in the mechanical properties of connective tissues. J Anat 131, 611–620.

Rafferty, N.S., and Scholz, D.L. (1985). Actin in polygonal arrays of microfilaments and sequestered actin bundles (SABs) in lens epithelial cells of rabbits and mice. Curr Eye Res 4, 713–718.

Rafferty, N.S., and Scholz, D.L. (1989). Comparative study of actin filament patterns in lens epithelial cells. Are these determined by the mechanisms of lens accommodation? Curr Eye Res 8, 569–579.

Rafferty, N.S., Scholz, D.L., Goldberg, M., and Lewyckyj, M. (1990). Immunocytochemical evidence for an actin-myosin system in lens epithelial cells. Exp Eye Res 51, 591–600.

Reilly, M.A. (2014). A quantitative geometric mechanics lens model: insights into the mechanisms of accommodation and presbyopia. Vision Res 103, 20–31.

Reilly, M.A., and Ravi, N. (2010). A geometric model of ocular accommodation. Vision Res 50, 330–336.

Scarcelli, G., Kim, P., and Yun, S.H. (2011). In vivo measurement of age-related stiffening in the crystalline lens by Brillouin optical microscopy. Biophys J 101, 1539–1545.

Schmut, O. (1978). The organization of tissues of the eye by different collagen types. Albrecht Von Graefes Arch Klin Exp Ophthalmol 207, 189–199.

Shi, Y., De Maria, A., Lubura, S., Sikic, H., and Bassnett, S. (2014). The penny pusher: a cellular model of lens growth. Invest Ophthalmol Vis Sci 56, 799–809.

Starodubtseva, M.N. (2011). Mechanical properties of cells and ageing. Ageing Res Rev 10, 16–25.

Sugiyama, Y., Akimoto, K., Robinson, M.L., Ohno, S., and Quinlan, R.A. (2009). A cell polarity protein aPKClambda is required for eye lens formation and growth. Dev Biol 336, 246–256.

Szczesny, S.E., and Elliott, D.M. (2014). Interfibrillar shear stress is the loading mechanism of collagen fibrils in tendon. Acta Biomater 10, 2582–2590.

Tsujii, A., Nakamura, N., and Horibe, S. (2017). Age-related changes in the knee meniscus. Knee 24, 1262–1270.

Upton, M.L., Gilchrist, C.L., Guilak, F., and Setton, L.A. (2008). Transfer of macroscale tissue strain to microscale cell regions in the deformed meniscus. Biophys J 95, 2116–2124.

Wang, K., Venetsanos, D.T., Wang, J., Augousti, A.T., and Pierscionek, B.K. (2017). The importance of parameter choice in modelling dynamics of the eye lens. Sci Rep 7, 16688.

Wanko, T., and Gavin, M.A. (1959). Electron microscope study of lens fibers. J Biophys Biochem Cytol 6, 97–102.

Willekens, B., and Vrensen, G. (1981). The three-dimensional organization of lens fibers in the rabbit. A scanning electron microscopic reinvestigation. Albrecht Von Graefes Arch Klin Exp Ophthalmol 216, 275–289.

Won, G.J., Fudge, D.S., and Choh, V. (2015). The effects of actomyosin disruptors on the mechanical integrity of the avian crystalline lens. Mol Vis 21, 98–109.

Zampighi, G.A., Eskandari, S., and Kreman, M. (2000). Epithelial organization of the mammalian lens. Exp Eye Res 71, 415–435.

Ziebarth, N.M., Wojcikiewicz, E.P., Manns, F., Moy, V.T., and Parel, J.M. (2007). Atomic force microscopy measurements of lens elasticity in monkey eyes. Mol Vis 13, 504–510.

